# The consequences of size-selective fishing mortality for larval production and sustainable yield in species with obligate male care

**DOI:** 10.1101/2020.03.24.006239

**Authors:** Holly K. Kindsvater, Kim Tallaksen Halvorsen, Tonje Knutsen Sørdalen, Suzanne H. Alonzo

## Abstract

Size-based harvest limits or gear regulations are often used to manage fishing mortality and ensure the spawning biomass of females is sufficiently protected. Yet, management interactions with species’ mating systems that affect fishery sustainability and yield are rarely considered. For species with obligate male care, it is possible that size-specific harvest of males will decrease larval production. In order to examine how size-based management practices interact with mating systems, we modeled fisheries of two species with obligate care of nests, corkwing wrasse (*Symphodus melops*, Labridae) and lingcod (*Ophiodon elongatus*, Hexigrammidae) under two management scenarios, a minimum size limit and a harvest slot limit. We simulated the population dynamics, larval production, and yield to the fishery under a range of fishing mortalities. We also modeled size-dependent male care to determine its interaction with management. In both species, the slot limit decreased yield by less than 12% (relative to minimum size limits) at low fishing mortalities; at higher mortalities, individuals rarely survived to outgrow the slot and spawning potential decreased substantially relative to unfished levels, similar to minimum size limits. Spawning potential decreased less when managed with a slot limit if we included a positive feedback between male size, care, and hatching success, but the benefit of implementing the slot depended both on the relative proportions of each sex selected by the fishery, and on our assumptions regarding male size and care. This work highlights that the effects of size- and sex-selective fisheries management can be nuanced and produce counter-intuitive results.

## Introduction

The sustainability of a fishery depends on the intensity of fishing pressure, the selectivity of the fishery for different sizes, ages, or sexes, and natural variation in population age and size structure arising from the species’ biology (Lowerre-Barbieri et al., 2017). Sustainable fisheries management aims to maximize yield to the fishery, either in numbers or in biomass, over the long term, by controlling total fishing mortality, the selectivity of the fishery, or both. To achieve this aim, management must also account for annual changes in population abundance and age structure due to fishing, natural environmental variation, and biological responses of the fished species to these processes (Halliday & Pinhorn, 2009; Zhou et al., 2010). For example, generation time, body size, and age at maturation are positively correlated with a stock’s risk of overexploitation and play a significant role in determining the stock-recruitment relationship (Hutchings & Kuparinen, 2017; Hutchings & Reynolds, 2004; Kindsvater, Mangel, Reynolds, & Dulvy, 2016).

The potential effects of fishing on mating system dynamics and recruitment success have rarely been incorporated into management, despite evidence fishing can affect these processes (Alonzo & Mangel, 2004; Kendall & Quinn, 2013; Rowe & Hutchings, 2003; Sørdalen et al., 2018). The effects of fishing on mating systems may be most important when fishing targets a specific size in sexually dimorphic species or in fisheries where management seeks to protect mature females from fishing (Alonzo & Mangel, 2004; Carroll & Lowerre-Barbieri, 2019; Kindsvater, Reynolds, Sadovy de Mitcheson, & Mangel, 2017; Sørdalen et al., 2018). In such cases, differential mortality between the sexes and altered sex ratios can increase the likelihood of egg or sperm limitation (Alonzo & Mangel 2004; Gosselin, Sainte-Marie, & Bernatchez, 2005; Hines et al., 2003). While the importance of large females to egg production and larval survival is well acknowledged (Arlinghaus, Matsumura, & Dieckmann, 2010; Birkeland & Dayton, 2005; Hixon, Johnson, & Sogard, 2014), the reproductive role of (large) males is often ignored (but see White, Cole, Cherr, Connon, & Brander, 2017). Males and sperm may also be limited (Alonzo *et al*. 2008; Sato 2012), body size can have positive paternal effects (Uusi-Heikkilä, Kuparinen, Wolter, Meinelt, & Arlinghaus, 2012), and theory predicts reduced reproductive rates when females are choosy but males with preferred phenotypes are scarce (Møller & Legendre, 2001).

The interaction between fishing and species with obligate male care, in which territorial males must defend eggs laid in nests to ensure eggs successfully hatch, has rarely been studied despite the fact that several commercially and recreationally important species display this behavior (Table 1; Halvorsen et al. 2017; King and Withler 2005; Sutter et al., 2012). In both cases, male care has presumably evolved to increase egg survival, and possibly to increase mating success via mate choice (Stiver & Alonzo, 2009). However, the relationship between care and selection on male and female size is complex, especially in species with alternative male reproductive tactics, where nesting male mating success (e.g., number of mates; paternity of the clutch he cares for) will also depend on the frequency of sneaking. For these species, size-selective fishing could decrease larval production by limiting the availability of nests, or the quality of male care, by selecting for smaller males. Smaller males may have smaller nests, fewer nests or provide less effective care (Cargnelli & Neff, 2006; Ingebrigt Uglem & Rosenqvist, 2002; Wiegmann & Baylis, 1995). Moreover, the most effective guarding males could be the largest, most aggressive males, so that the traits that correlate with the most effective parental care could also increase vulnerability to fishing (Andersen, Marty, & Arlinghaus, 2018; Sutter et al., 2012).

**Table 1.** Fished species with male care.

| Species | Larger sex | Alternative male tactic | Hermaphroditism | Habitat | Fishery | Management includes slot limit (for some populations) |
| --- | --- | --- | --- | --- | --- | --- |
| Ballan wrasse (*Labrus bergylta*, Labridae) | Male | No | Protogynous | Marine | Commercial and recreational | Yes |
| Cuckoo wrasse (*Labrus mixtus*, Labridae) | Male | Yes | Protogynous | Marine | Commercial | Yes |
| Corkwing wrasse (*Symphodus melops*, Labridae) | Male | Yes | No | Marine | Commercial | Yes |
| Black bream (*Spondyllosoma cantharus*, Sparidae) | Male | No | Protogynous | Marine | Recreational | No |
| Bluegill sunfish (*Lepomis macrochirus*, Centrarchidae) | Male | Yes | No | Fresh water | Recreational | No |
| Largemouth bass (*Micropterus salmoides*, Centrarchidae) | Monomorphic | No | No | Fresh water | Recreational | Yes |
| Smallmouth bass (*Micropterus dolomieu*, Centrarchidae) | Monomorphic | No | No | Fresh water | Recreational | Yes |
| Atlantic wolffish (*Anarhichas lupus*, Anarhichadidae) | Monomorphic | No | No | Marine | Commercial | No |
| Butterfly peacock bass (*Cichla ocellaris*, Cichlidae)* | Monomorphic | No | No | Fresh water | Recreational | No |
| Lingcod (*Ophiodon elongatus*, Hexagrammidae) | Female | No | No | Marine | Commercial and recreational | Yes |
| Lumpfish (*Cyclopterus lumpus*, Cyclopteridae) | Female | No | No | Marine | Commercial | No |

\*Note: Butterfly peacock bass has biparental care.

Despite growing recognition of the importance of maintaining diverse size structure in fished populations, there is little theory predicting how size-selective fishing will interact with reproductive behavior and influence population productivity. Conventional wisdom suggests size-selective management (e.g., minimum size limits), designed to allow individuals to mature and reproduce successfully before being fished, will ensure sustainable yields in the long term (Birkeland, & Dayton, 2005; Froese, 2004). More recently, protection of the largest, oldest individuals has been recognized as a desirable management outcome if large individuals play important roles ecologically or contribute high quality larvae to the population (Ahrens, Allen, Walters, & Arlinghaus, 2020; Hixon, Johnson, & Sogard, 2014). A maximum size limit, in addition to a minimum size limit, often referred to as a harvest slot, has been suggested to prevent sex-selective harvest when one sex grows faster than the other (Halvorsen et al., 2016; Morson, Munroe, Harner, & Marshall, 2017). Slot limits have been increasingly implemented in recreational fisheries as a means of ensuring that fishing’s effects on stock size, age, or sex structure are balanced, or to maintain a suitable number of large fish available to anglers (Gwinn et al., 2015). Slot limits have been advocated in fisheries for species in which mating success and egg survival also depend on body size as in species with obligate paternal care (Halvorsen et al., 2017). However, when considering the effectiveness of a minimum size limit vs. a slot limit, it can be unclear how each management action will interact with the species’ mating system and sex-specific growth differences to affect fishery sustainability and yield. For example, in species with paternal care, it is unclear whether it is more important for management actions to protect females (typically the gamete-limiting sex) or males to ensure a sustainable larval supply.

To explore these alternatives, we use simulations of age-, size- and sex-structured population dynamics to understand how fishing will interact with natural variation in male and female size structure. We focus on comparing the effects of different size-selective management scenarios in species with obligate male care, but different mating systems and different sex-specific growth patterns. With a model of age-, size- and sex-structured population dynamics, we compare the effects of a fishery selecting predominantly the large nesting males of a species with multiple male phenotypes, the corkwing wrasse (*Symphodus melops*, Labridae), with a fishery for a species where females are the larger sex, and both males and females are caught, lingcod (*Ophiodon elongatus*, Hexigrammidae). Both of these species have obligate male care of eggs laid in benthic nests. We specifically evaluate the consequences of alternative management tactics (a minimum size limit vs. a slot limit) on the production of eggs and recruits by calculating the Spawning Potential Ratio (SPR) and yield in numbers. By tracking male and female numbers and sizes in the model, we are able to quantify the effects of management on both egg production and the availability of paternal care. In comparing these species, we aim to gain general insights into the way that species biology and fishery selectivity interact.

As fishing removes males of a specific size, the average size of a nesting male is expected to decrease, because fewer males will survive to the largest age and size class. Whether fishing affects larval production in species with male care depends on the consequences of size- and sex-selective fishing for the egg density in each nest, and the extent to which egg survivorship depends on density. In fishes, male care is often regarded as shareable among eggs, meaning that additional eggs do not decrease the quality of care, although there is some evidence for density-dependent egg survival within a nest (Klug, Lindstrom, & St. Mary, 2006). In addition, male size has shown to be positively correlated to the intensity of care and nest survival (Suski & Ridgway, 2007; Wiegmann & Baylis, 1995). Here, we develop a new metric of the availability of care *per egg* and quantify how that is expected to change with the impacts of fishing on nest size and number, as well as on egg production. This is the first theoretical investigation of the interplay between fishing and larval production in species with alternative mating tactics and obligate male care.

## Study species

We modelled two contrasting fisheries for species with obligate male care: the commercial fishery for live corkwing wrasse in southern and western Norway and the recreational fishery for lingcod in western North America. These are among the two best-studied marine fisheries for species with parental care, and each fishery has a history of management that includes minimum size limits and slot limits. Both corkwing wrasse and lingcod males will defend eggs laid by one or more females in benthic habitat for a period of 6 to 8 weeks, depending on water temperature.

### Corkwing wrasse

The corkwing wrasse has two male mating tactics, with large nesting males providing care for eggs and sneaker males that steal fertilizations during spawning events (Potts, 1974; I. Uglem, Rosenqvist, & Wasslavik, 2000). Male and female corkwing wrasse have different growth and maturation rates, as do the two male life-history pathways (territorial nesting males and sneakers (Halvorsen et al., 2016; Uglem, Rosenqvist, & Wasslavik, 2000). Sneaker males mature after one year. Nesting males and females mature between two and three years, and can live for up to nine years.

The parameters for growth and maturation rates in our model were estimated from the data published in Halvorsen et al. (2016), combined with more recent samples collected in 2017. Growth and maturation rates as a function of age are plotted in Figure 1 (a,c); functions are in Table 2; Parameter estimates are in Table 3. We assume mass-at-age *W*(*a*) is a cubic function of length, and we estimate egg production as a function of mass following the estimates in Chalaris (2011). Like many batch spawners, corkwing wrasse are income breeders, meaning that food availability affects reproductive output (number of eggs spawned) over short time scales. Very little is known about how many times the females spawn each season, although it is more than once, and polyandry is common (Stiver et al., 2018; Ingebrigt, Uglem & Rosenqvist, 2002). While we do not have direct information on natural mortality rates for each life-history type of corkwing wrasse, estimates of total mortality (natural mortality and fishing mortality) can be estimated from the frequency of each age class by life-history type in the catch data (Halvorsen *unpublished data*). The commercial fishery for live corkwing wrasse has grown rapidly since 2010 as the wrasse are used in louse biocontrol in salmon aquaculture (Blanco Gonzalez et al., 2019; Halvorsen et al., 2017). The fishery is size-selective, as the largest individuals fare better in salmon pens, and since nesting males grow faster and mature later than females and sneaker males, their life history is less protected by the current minimum size regulations. To account for this, a slot harvest limit – where only fish between 120 and 170 mm can be retained – has been proposed to protect the largest males in response to scientific concerns about the effects of fishing on the mating system (Fig 1a; Halvorsen et al., 2017). In England, corkwing wrasse is managed by slot sizes in two districts (IFCA 2020).

**Figure 1.**
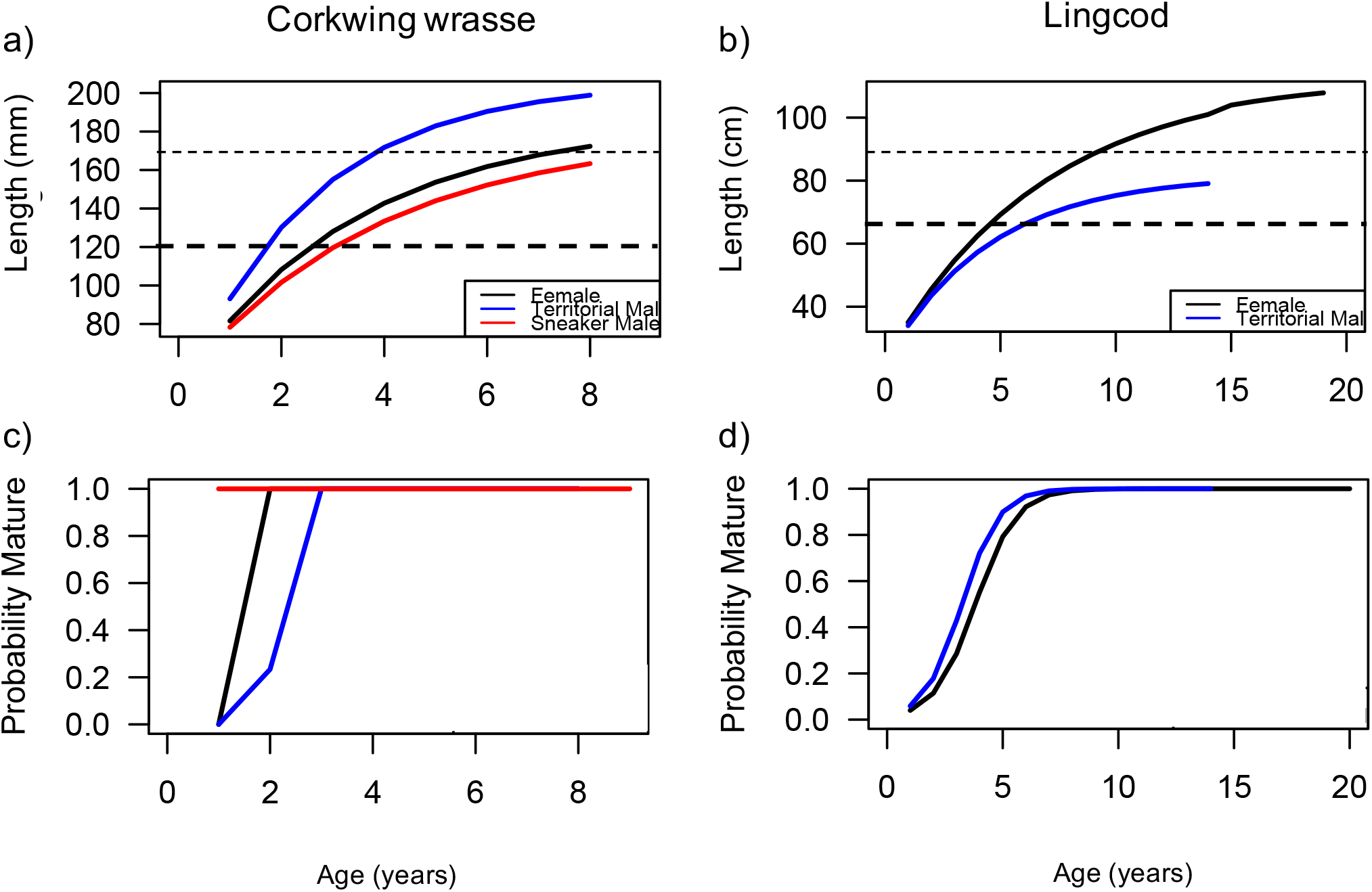
Length-at-age functions and age-specific maturation rates for corkwing wrasse (left column) and lingcod (right column). For the wrasse, estimates of growth (panel a) and maturation rates (panel b) are available for each life-history pathway in the western Norway stock (females: black line; nesting males: blue line; sneaker males: red line). For lingcod, sexspecific estimates (females: black; males: blue) are available from stocks in Washington State. Note female lingcod are the larger sex. Parameters for each function are given in Table 3.

**Table 2.**
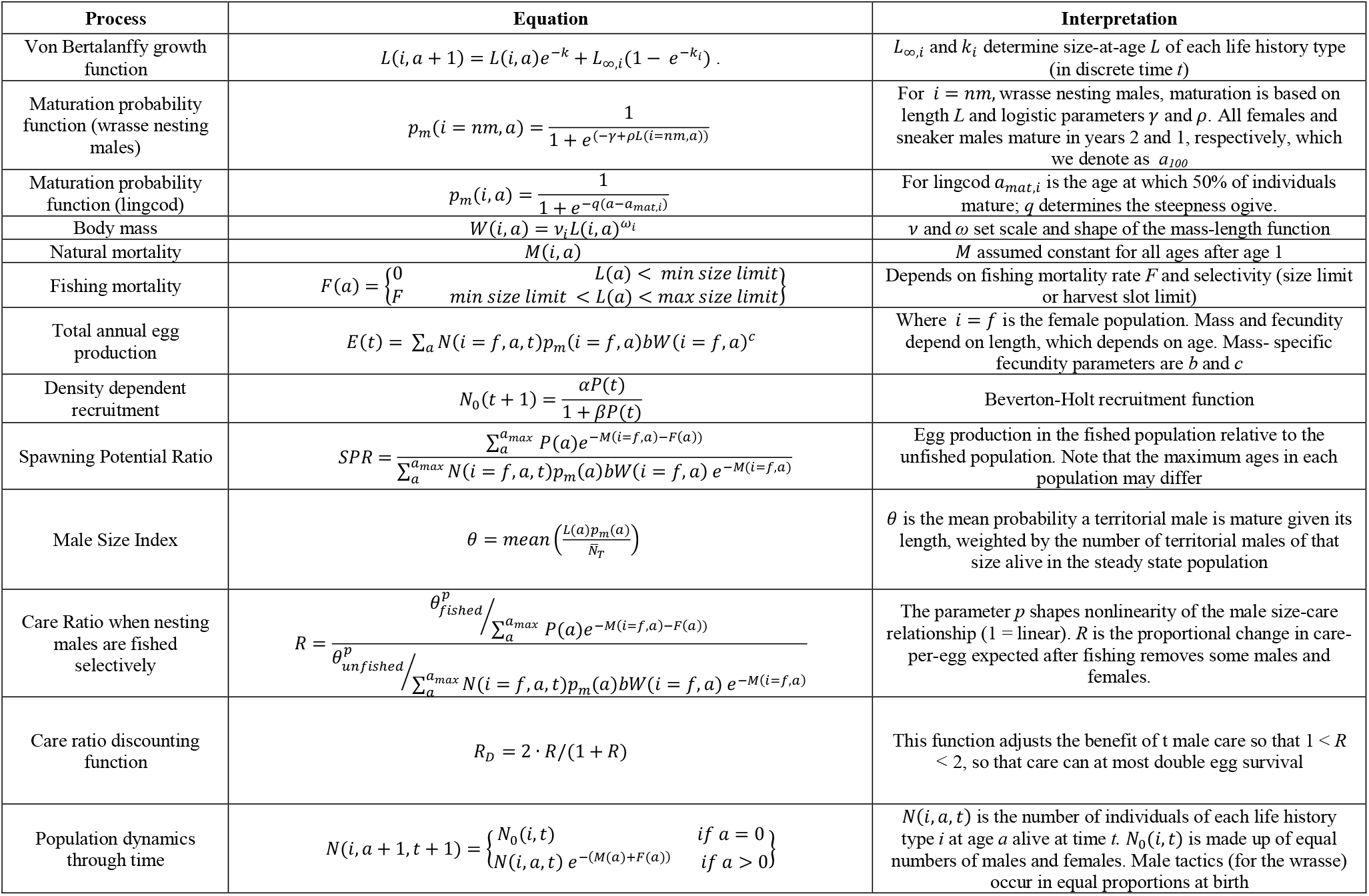
Population model functions and interpretation

| Process | Equation | Interpretation |
| --- | --- | --- |
| Von Bertalanffy growth function | $L(i, a + 1) = L(i, a)e^{-k} + L_{\infty,i}(1 - e^{-k_i})$ | $L_{\infty,i}$ and $k_i$ determine size-at-age $L$ of each life history type (in discrete time $t$) |
| Maturation probability function (wrasse nesting males) | $p_m(i = nm, a) = \frac{1}{1 + e^{(-\gamma + \rho L(i=nm,a))}}$ | For $i = nm$, wrasse nesting males, maturation is based on length $L$ and logistic parameters $\gamma$ and $\rho$. All females and sneaker males mature in years 2 and 1, respectively, which we denote as $a_{100}$ |
| Maturation probability function (lingcod) | $p_m(i, a) = \frac{1}{1 + e^{-q(a - a_{mat,i})}}$ | For lingcod $a_{mat,i}$ is the age at which 50% of individuals mature; $q$ determines the steepness ogive. |
| Body mass | $W(i, a) = v_i L(i, a)^{\omega_i}$ | $v$ and $\omega$ set scale and shape of the mass-length function |
| Natural mortality | $M(i, a)$ | $M$ assumed constant for all ages after age 1 |
| Fishing mortality | $F(a) = \begin{cases} 0 & L(a) < \text{min size limit} \\ F & \text{min size limit} < L(a) < \text{max size limit} \end{cases}$ | Depends on fishing mortality rate $F$ and selectivity (size limit or harvest slot limit) |
| Total annual egg production | $E(t) = \sum_a N(i = f, a, t) p_m(i = f, a) b W(i = f, a)^c$ | Where $i = f$ is the female population. Mass and fecundity depend on length, which depends on age. Mass- specific fecundity parameters are $b$ and $c$ |
| Density dependent recruitment | $N_0(t + 1) = \frac{\alpha P(t)}{1 + \beta P(t)}$ | Beverton-Holt recruitment function |
| Spawning Potential Ratio | $SPR = \frac{\sum_a^{a_{max}} P(a) e^{-M(i=f,a)-F(a)}}{\sum_a^{a_{max}} N(i = f, a, t) p_m(a) b W(i = f, a) e^{-M(i=f,a)}}$ | Egg production in the fished population relative to the unfished population. Note that the maximum ages in each population may differ |
| Male Size Index | $\theta = \text{mean}\left(\frac{L(a)p_m(a)}{N_T}\right)$ | $\theta$ is the mean probability a territorial male is mature given its length, weighted by the number of territorial males of that size alive in the steady state population |
| Care Ratio when nesting males are fished selectively | $R = \frac{\theta_{fished}^p / \sum_a^{a_{max}} P(a) e^{-M(i=f,a)-F(a)}}{\theta_{unfished}^p / \sum_a^{a_{max}} N(i = f, a, t) p_m(a) b W(i = f, a) e^{-M(i=f,a)}}$ | The parameter $p$ shapes nonlinearity of the male size-care relationship ($1 = \text{linear}$). $R$ is the proportional change in care-per-egg expected after fishing removes some males and females. |
| Care ratio discounting function | $R_D = 2 \cdot R / (1 + R)$ | This function adjusts the benefit of $t$ male care so that $1 < R < 2$, so that care can at most double egg survival |
| Population dynamics through time | $N(i, a + 1, t + 1) = \begin{cases} N_0(i, t) & \text{if } a = 0 \\ N(i, a, t) e^{-(M(a)+F(a))} & \text{if } a > 0 \end{cases}$ | $N(i, a, t)$ is the number of individuals of each life history type $i$ at age $a$ alive at time $t$. $N_0(i, t)$ is made up of equal numbers of males and females. Male tactics (for the wrasse) occur in equal proportions at birth |

**Table 3.** Simulation parameters

|  |  |  |  |  |  |  |
| --- | --- | --- | --- | --- | --- | --- |
| $N$ | Initial population size (assumed 50:50 sex ratio, 25% of males assumed to be sneakers) | 1000 | | | | |
| $T_{max}$ | Number of years in our population simulation. | 200 | | | | |
| $T_{fishin\ g}$ | Time when fishing mortality begins | 100 | | | | |
|  |  | Corkwing wrasse parameters |  |  | Lingcod parameters |  |
|  |  | Females | Territorial males | Sneaker males | Females | Territorial males |
| $M(a)$ | Natural mortality rate (assumed constant) | 0.54 | 0.54 | 0.40 | 0.18 | 0.32 |
| $A_{max}$ | Maximum age (years) | 8 | 8 | 9 | 20 | 14 |
| $L_{inf}$ | Asymptotic size in von Bertalanffy growth function | 177.5 mm | 199.72 mm | 180.8 mm | 112.8 cm | 81.87 cm |
| $k$ | Growth coefficient in von Bertalanffy growth function | 0.2972 | 0.3994 | 0.2625 | 0.145 | 0.223 |
| $L$ | Length-at-age; $L(1)$ is reported here as the initial size at recruitment | 81.6 mm | 93.1 mm | 78.3 mm | 35 cm | 34 cm |
| $\nu$ | Scale coefficient relating length to body mass (in kg) | $6.6\times10^{-7}$ | $9.3\times10^{-7}$ | $8.5\times10^{-7}$ | $1.7\times10^{-3}$ | $4.0\times10^{-3}$ |
| $\omega$ | Shape exponent relating length to body mass | 3.176 | 3.089 | 3.115 | 3.40 | 3.21 |
| $a_{100}$ | Age at maturation (wrasse females/sneaker males) | 2 | - | 1 | - | - |
| $\gamma$ | Coefficient of length-based maturation function for wrasse nesting males | - | 18.5836 | - | - | - |
| $\rho$ | Shape coefficient of length-based maturation function for wrasse nesting males | - | 0.13361 | - | - | - |
| $a_{mat}$ | Age when 50% of lingcod are mature | - | - | - | 1.129 | 1.124 |
| $q$ | Steepness of age-based maturation function for lingcod | - | - | - | 3.814 | 3.233 |
| $b$ | Mass-specific fecundity coefficient (mass in kg) | $2.95\times10^5$ | - | - | $2.82\times10^{-4}$ | - |
| $c$ | Mass-specific fecundity exponent | 1 | | | 3.0011 | - |
| Recruitment parameters |  |  |  |  |  |  |
| $\alpha$ | Slope near the origin of the Beverton-Holt recruitment function | 0.05 (0.03-0.07) | | | 0.1 (0.8-0.11) | |
| $\beta$ | Parameter of the Beverton-Holt recruitment function | $8\times10^{-6}$ ($8\times10^{-5}$) | | | $1\times10^{-5}$ ($8\times10^{-4}$) | |
| Exploitation parameters |  |  |  |  |  |  |
| - | Minimum size limit (mm) | 120 |  |  | 26 inches (66.04 cm) |  |
| - | Maximum size limit (slot) | 170 |  |  | 36 inches (91.44 cm) |  |
| $F$ | Fishing mortality rate | Ranged from 0 to 0.7 | | | | |

### Lingcod

We compared the effect of fisheries management on the wrasse system with lingcod, which has a contrasting life history (Figure 1, b,d). The lingcod is the only extant member of the family Hexigrammidae and is native to western North America. Lingcod are large, long-lived fish, although the ages of maturity of the nesting male wrasse and lingcod are both between two and three years (Cass, Beamish, McFarlane, & Canada. Department of Fisheries and Oceans., 1990; King & Withler, 2005). Lingcod females are capable of growing to over two meters long and they live, on average, six years longer than males, with lower natural mortality and later.

Estimates of growth, maturation, and mortality rates, as well as estimates of mass-specific egg production by females, and initial size at recruitment, were reported in Jagielo and Wallace (2005), an international stock assessment that encompassed populations from California to the Salish Sea. While there is undoubtedly some regional variation in growth and body size, these growth and maturation values are representative of the species and are useful when examining the effect of the contrasting life histories and mating systems of lingcod and corkwing wrasse (Figure 1). There are two notable differences in lingcod growth patterns and mating system that differ from the corkwing wrasse. First, female lingcod are the larger sex, and spawn one clutch with only one male each season (total spawning). Second, while smaller males are known to steal fertilizations by sneaking during spawning events, it is likely that male mating tactics change ontogenetically as males grow, and are not separate life history pathways (King and Withler 2005).

Lingcod have been fished commercially and recreationally for decades and were considered to be overfished by the late 20th century due to intense fishing in the 1980s and 1990s (Haltuch et al., 2017; Jagielo & Wallace, 2005). Management has varied among Alaska, British Columbia, Washington, Oregon, and California according to the history of the fishery and natural latitudinal variation in growth and body size. Current management practices are a mix of area closures, bag limits, minimum-size limits, and slot limits in different parts of the range. In Washington State, recreational fishers are regulated by a slot limit of 60-91 cm (Fig 1b). Due to the implementation of precautionary management measures in recent years, lingcod populations are rebuilding and appear to be stable throughout much of the range (Haltuch et al., 2017).

## Methods

We developed a deterministic model of equilibrium population dynamics (Table 2), which is closely related to the size-and age-structured model commonly used in fisheries stock assessments (Mangel 2006). However, our model differs in several important ways. First, we allowed for the sexes to have different growth rates. We also addressed the two growth patterns of the different male life history strategies (nesting male and sneaker) for the corkwing wrasse population. Second, we modeled differences in natural mortality and fishery susceptibility for each life-history type. Third, we considered the relationship between the male size structure, paternal care, and larval production. Figure 2 shows a general schematic of the model. With this model, we can evaluate the potential consequences of fishing for population productivity, given different assumptions regarding how male size structure affects the availability of paternal care in nests and the resulting consequences for the production of larvae.

**Figure 2.**
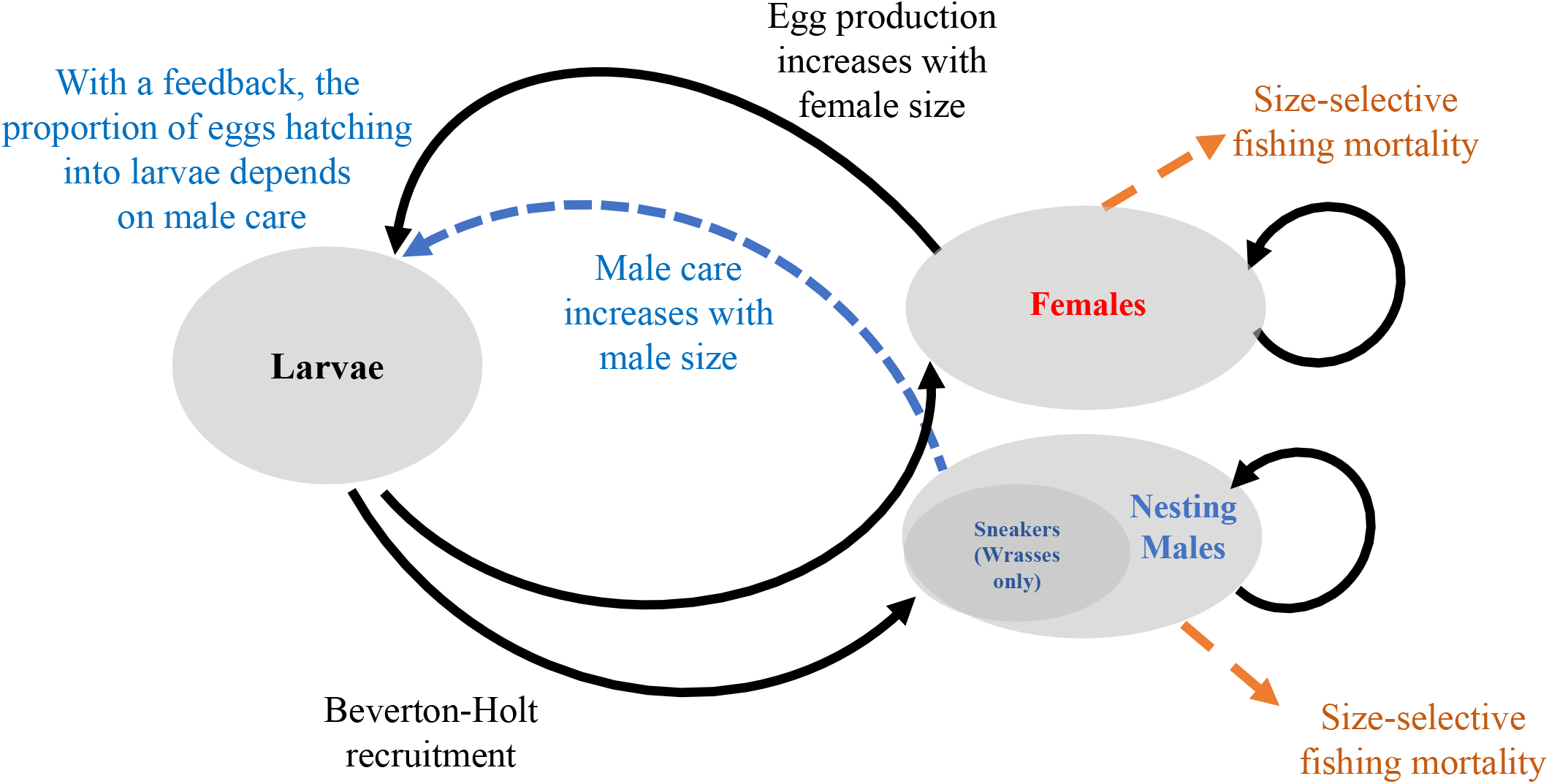
Generic schematic of the population model and where the reproductive behavior operates. The model assumes that recruitment to the adult population happens at after one year of age for both species. After that, they experience constant annual mortality, which varies according to sex and male tactic. Maturation rates for each sex and life history vary according to age and/or size, depending on how they were estimated empirically (Table 3). Female fecundity depends on her mass, which is a function of her age in this model. In the scenarios where we included an effect (feedback) of male care, the probability an egg hatched and becomes a planktonic larva is proportional to the Care Ratio R, based on the average size of territorial mature males, weighted by their frequency in a stable age distribution under fishing, relative to their frequency in the unfished population (Table 2).

### Male size and paternal care

We assume the effectiveness of paternal care in a nest – which is essential for the production of viable larvae in both species – increases with the size of nesting males in both corkwing wrasse and lingcod. In both species, larger males have larger nests and nest multiple times in a season or defend multiple nests at once (King & Withler, 2005b; Potts, 1985; Ingebrigt Uglem & Rosenqvist, 2002). It is also possible that large males are more effective at nest defense or fanning than small nesting males (Coleman & Fisher, 1991; Wiegmann & Baylis, 1995; Lissåker & Kvarnemo, 2006). However, the exact relationship between egg number and larval numbers as a function of male size and nest size is unknown for these and most other species. Per-capita survival of eggs in the nest could be a decreasing function of egg density and nest size (i.e., larval production could be a density-dependent function of egg number due to crowding). Both males and females may be adapted to avoid spawning under conditions of nest crowding, but if the number of nests is limited, they might choose to spawn anyway. As there are a range of possibilities, as a starting point, we chose a simple relationship between male size and available nest care to represent both male nest size and number of nests. We developed a Male Size Index, which we represent as *θ*, representing the probability that a territorial male of a given length is mature, and the average length of territorial males, weighted by the abundance of each age class (Table 2). We modeled available nest care as a power function, such that care *C* = *θ^p^*. We varied + from one (linear) to 1.25 (slightly concave up). We used the nest care function to calculate the per-egg availability of care in a nest as the ratio of available care *C* to total egg production each year. Both care availability and egg production decreased with fishing, but the selectivity of fishing pressure for each sex determined whether care-per-egg decreased or increased. We characterize this change as the Care Ratio, which we represent as *R:* the care-per-egg in the fished population, relative to the care-per-egg in the unfished population (Table 2).

Using this new metric *R*, we evaluated the consequences of an effect of fishing on larval production (the total number of eggs that hatch successfully each year) due to a change in care-per-egg. We assumed larval hatching success changed proportionally according to the Care Ratio in this subset of our analyses. The way we modeled hatching success depended on whether care-per-egg changed with fishing mortality relative to the unfished population. If available care decreased (i.e., *R* < 1), we assumed this also decreased larval hatching success by the same fraction. However, we observed that when fishing on females was more intense than males, as in lingcod, it was possible for care-per-egg to increase (*R* > 1). While we think that increased male care could potentially improve per-egg survival, under some fishing scenarios *R* increased dramatically. As the relationship between care and per-offspring survival generally asymptotes, we added a function that meant increased care saturated at two (so it doubled per-egg hatching success; Table 2). In this way, we estimated the indirect effect of changes in per-egg availability of care after fishing removed some males and females. Our goal was to compare the net effect of these changes to male and female demography on larval production and yield to the fishery, with and without the potential feedback between care availability and hatching success. In every scenario, we calculated expected yield and larval production, sometimes referred to as spawning potential. The latter is correlated with a population’s capacity to buffer stochastic environmental variation (Kindsvater et al., 2016; O’Farrell & Botsford, 2005).

### Recruitment and population dynamics of both species

Births and deaths in our simulated population were determined by the demographic composition of males and females. The maturation rate and mortality rate of each sex of each species determines the number of mature individuals in each age class (*a*) alive in each time step, *N*(*t*), if they are mature *p_m_*(*a*), and the relationship between fecundity and body mass *W*(*a*). Egg production *E*(*t*) depends on the number and size of mature females, and the mass-fecundity relationship for each species (Table 2; parameters are given in Table 3). In scenarios with no effect of care availability, egg production and larval production *P*(*t*) were perfectly correlated, *E*(*t*) = *P*(*t*). As illustrated in Figure 2, with a feedback incorporating care availability, larval production *P*(*t*) was adjusted according the Care Ratio, so that *P*(*t*) = *RE*(*t*). For all scenarios, we assumed the population dynamics are regulated by density-dependence in recruitment to the year-one age class. In other words, larval survival from hatching to recruitment depends on larval density, but adult survival and growth are independent of density. We assumed the recruitment function followed the Beverton-Holt equation (Mangel, 2006). The Beverton-Holt relationship specifies the maximum probability that a larva survives to recruitment *α* (i.e., the productivity of the population at low density) and a metric of the strength of density dependence *β*. These parameters arise from specific environmental conditions that are difficult to measure, so they are often estimated from stock-recruitment relationships that relate recruitment to spawning stock biomass (Mangel, 2006). In species lacking stock-recruitment relationships, such as the corkwing wrasse, we can characterize the effects of fishing on a population relative to an arbitrary unfished population. For comparison, we completed the same analysis on lingcod. Importantly, as long as recruitment overfishing is not occurring, our assumptions regarding density-dependent recruitment (i.e., the Beverton-Holt stock recruitment curve) do not have a strong effect on relative yield and relative spawning potential (sometimes called the Spawning Potential Ratio, SPR). To make sure this assumption is valid for the results presented here, we used a range of values for *α* and *β* and checked that they did not influence on our conclusions (Ranges of the recruitment parameters are in Table 3).

After individuals recruit to the population model after 1 year at size *L_R_* (in cm), they grow and reproduce each year according to the specific growth, maturation, and mortality rates reported for each species (Table 3, Figure 2). Individuals live to at most *A_max_* years. Given the balance of birth and death rates, the population will equilibrate at a steady state population biomass. We checked that our simulated populations all reached this equilibrium within 100 years, at which time we “fished” our populations and made sure the fished population reached a new steady state. The selectivity of fishing mortality varied according to the management scenario we simulated. We evaluated a range of fishing mortality coefficients (*F* in Tables 2 and 3) for each scenario, which were chosen because they span the range from very low fishing mortality to quite intense harvest that is likely to lead to overfishing. As a reference, in our model *F_40_* = (when spawning biomass is reduced to 40% of unfished levels) was near *F* = 0.7 for corkwing wrasse and *F* = 0.5 for lingcod, depending on the management scenario. For each species, we compared a minimum size limit with a slot size limit, based on size limits currently used in management. We also compared scenarios with and without a feedback between care and larval production. This factorial comparison allowed us to tease apart the effects of management on the indirect effects of fishing on yield and larval production. Doing this for both species revealed the interaction between the details of each life history and management decisions.

For each factorial combination, we calculated yield to the fishery in numbers of fish, rather than biomass. To evaluate spawning potential, we compared the lifetime egg production of individuals in the fished population to those in the unfished, which is a method of calculating the Spawning Potential Ratio (SPR; Kindsvater et al., 2016). This reference point is commonly used to understand the recovery potential of a population, and conversely, its risk of overexploitation (O’Farrell & Botsford, 2005). Finally, we calculated the Care Ratio *R:* the care-per-egg available in the fished population, relative to the unfished population (Table 2).

## Results

### Corkwing wrasse

Figure 3 shows that implementing the harvest slot limit led to a small decrease in yield to the wrasse fishery. A decrease in yield after the implementation of the slot is expected, because the largest individuals are protected from fishing. Natural mortality is high enough in corkwing wrasse that very few individuals in our model survived to grow beyond the maximum size limit, especially under strong fishing mortality. Therefore, the percent change in yield between the minimum size limit and the slot scenario was greatest at low fishing mortality (Fig. 3b). This implies that for species with high natural mortality, the benefits of a slot limit will be countered by any corresponding change in fishing mortality on size classes within the slot. In other words, if the harvest rate of size classes within the slot increases as fishers respond to its implementation, it is possible the benefits of the slot will be undetectable.

**Figure 3.**
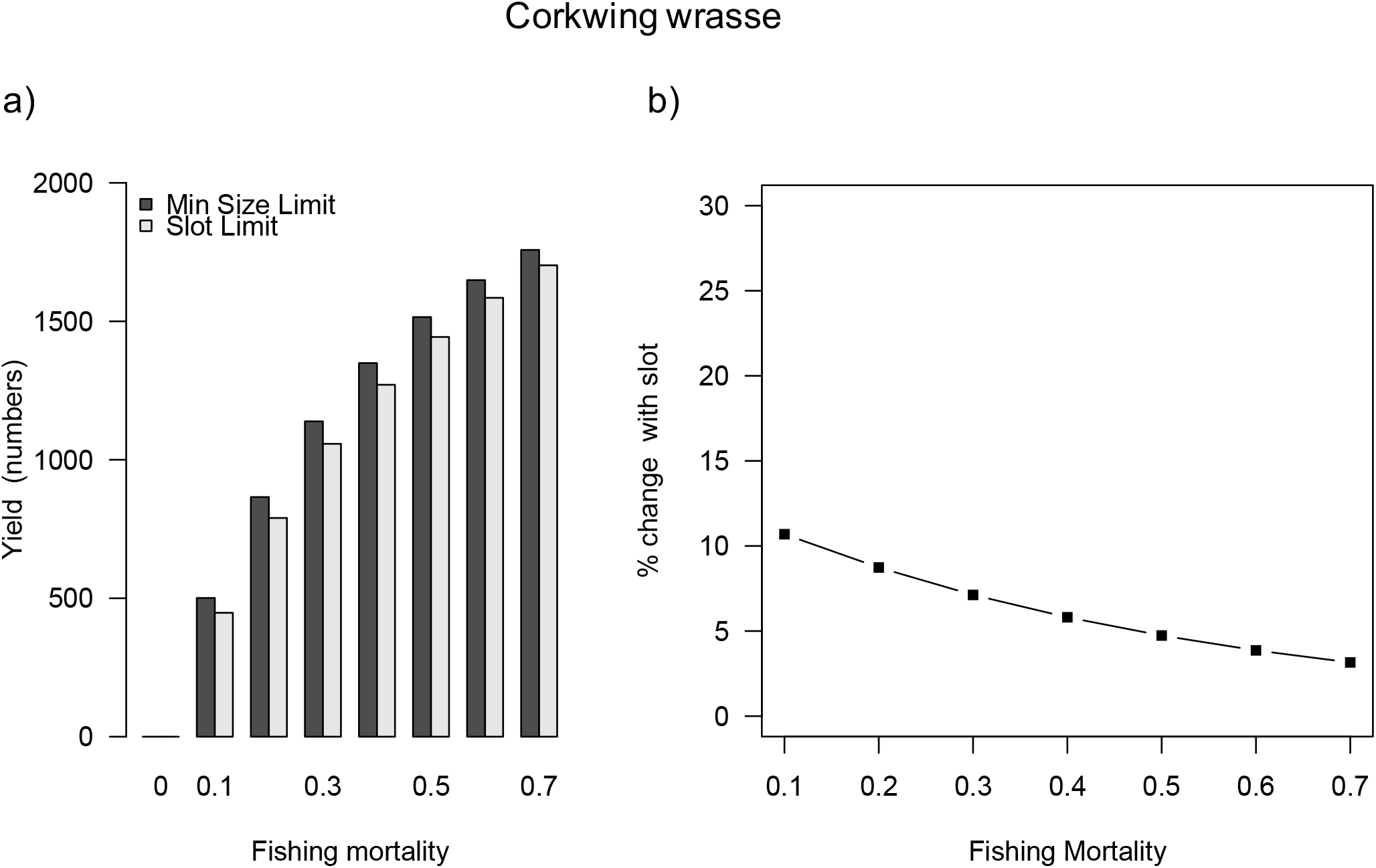
Yield of corkwing wrasse, in numbers of fish (panel a), for the minimum size limit (dark bars) and the slot limit (pale bars). We calculate the percent difference in yield under the two management scenarios (panel b). Yield is always greater under the minimum size limit, but that difference narrows as the population becomes overfished.

In our model, yield did not change in the scenarios including the feedback between male size structure and larval production. While the slot limit increased care capacity by decreasing the number of large males captured, the minimum size limit was sufficient to protect egg production enough that recruitment to the adult population (at age one) was not affected. This is a consequence of our use of a deterministic Beverton-Holt recruitment function. Because stock recruitment relationships vary according to inter-annual environmental fluctuations, the result that yield is stable under high size-selective fishing mortality is an oversimplification of reality and could lead to overconfidence in a fishery’s sustainability. Therefore, we focus on the Spawning Potential Ratio (a proxy for larval production) as an indicator of the ability of the population to buffer environmental stochasticity.

The dashed lines in each panel of Figure 4 show the Care Ratio *R* (small red circles), which reflects the care capacity of the population and the SPR (purple squares and blue circles) with and without feedbacks. As expected, with fishing, the spawning potential is lower when egg survival to the larval stage decreases with the availability of care, because of the truncated size structure of males. In other words, fishing has a stronger negative effect on larval supply when we assume there is an effect of care capacity (which depends on male size) on larval production. Implementing a harvest slot mitigates this effect slightly (compare Fig. 4a and 4b). Additionally, the efficacy of the slot in increasing larval supply depends on the shape parameter *p*, which determines nonlinearity of the relationship between male size and care capacity. At higher values of *p* (1 < *p* < 2), protecting the largest males with a slot limit has an outsized effect on the population’s care capacity and increases the SPR (with the feedback; purple squares in Fig 4b), especially at low levels of fishing mortality when the slot is most effective (not shown).

**Figure 4.**
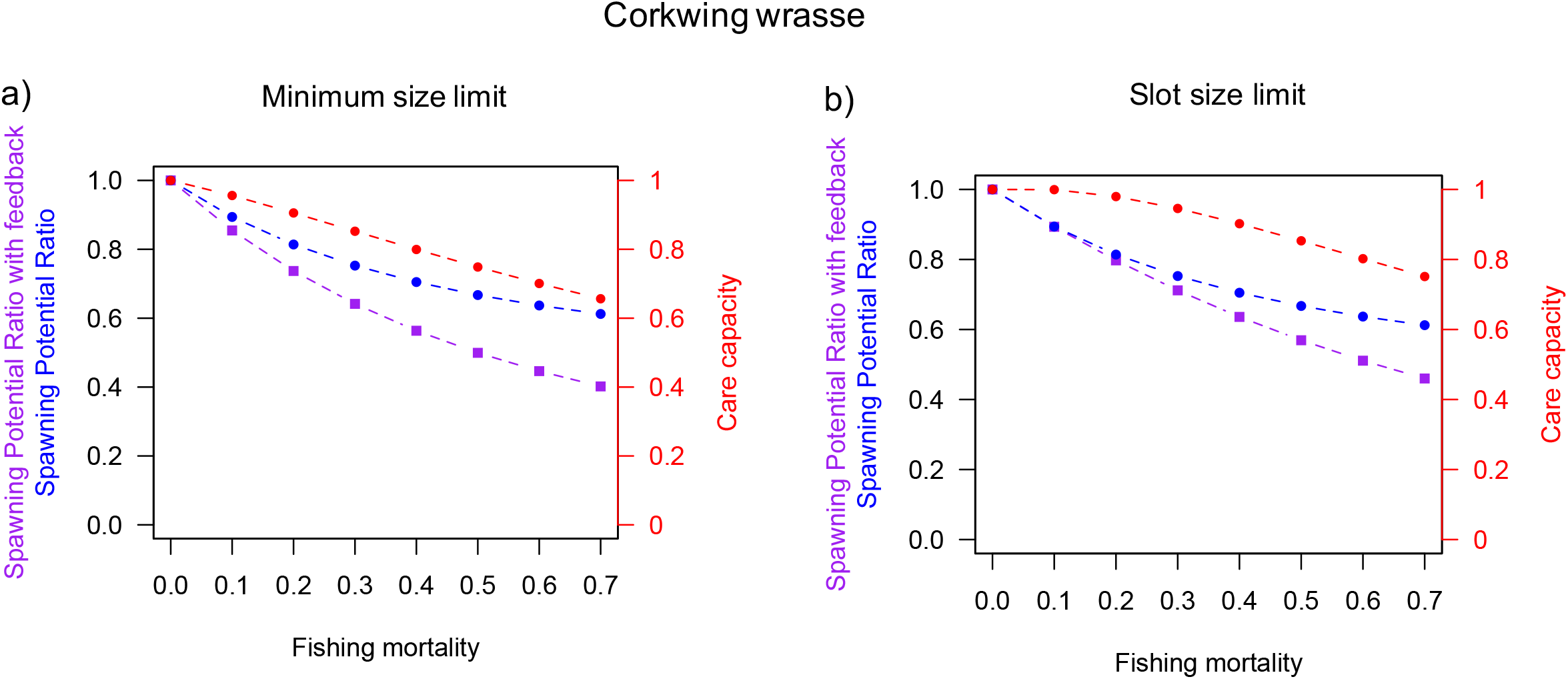
Spawning potential and care capacity for each of the factorial combinations of corkwing wrasse simulations. We modeled spawning potential of fisheries with a minimum size limit (panel a) or a slot limit (panel b), each with (purple-dashed squares) and without (blue-dashed circles) a feedback between care and larval production, across a range of fishing mortalities. Red-dashed circles represent the Care Ratio, R.

### Lingcod

As in the wrasse example, the yield to the fishery (in number of fish) decreased with the implementation of a harvest slot limit (Fig. 5), but did not vary with an effect of care capacity on larval production. Again, we attribute this stability in catch to our choice of the Beverton-Holt stock recruitment function and the fact the minimum size limit is sufficient to prevent recruitment overfishing in our deterministic model. As in the wrasse, yield differences arising from the two management scenarios was greatest at low fishing mortality. At high rates of fishing mortality, yield did not differ substantially between the minimum size and slot limit scenarios (Fig. 5), as few individuals survived to outgrow the maximum size limit.

**Figure 5.**
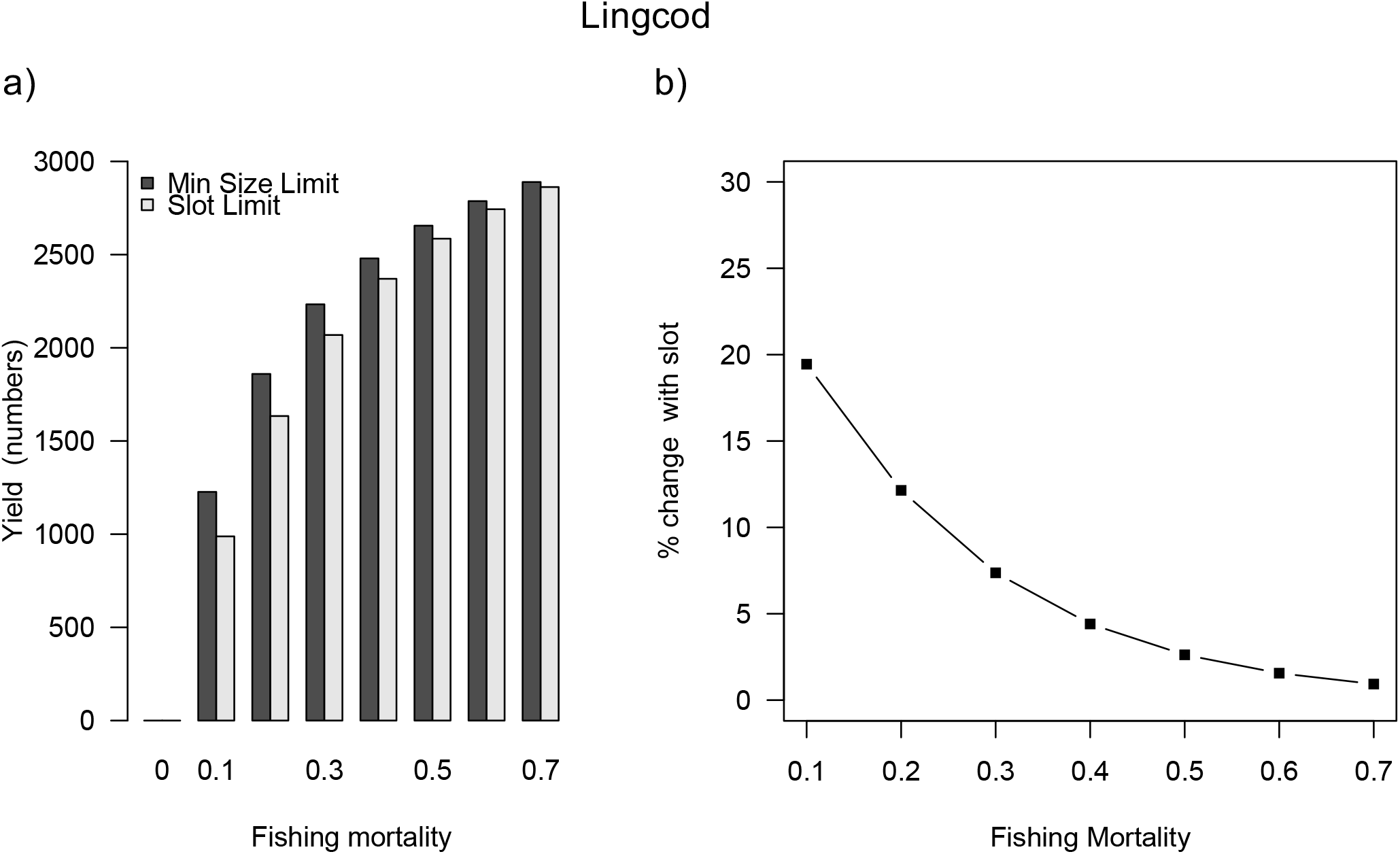
Yield of lingcod, in numbers of fish (panel a), for the minimum size limit (dark bars) and the slot limit (pale bars). We calculate the percent difference in yield under the two management scenarios (panel b). As in Fig. 3, yield is always greater under the minimum size limit, but that difference narrows as the population becomes overfished.

Lingcod females are larger than males, and the main effect of fishing was to strongly reduce the spawning potential of females (Fig. 6, blue circles). Including a feedback with care capacity mitigated this decrease in SPR (Fig. 6, purple squares). This pattern held in both management scenarios. This is because in lingcod, the availability of care increased *above* unfished levels after fishing, because egg production decreased faster than nests. The increase in care-per-egg is reflected in a Care Ratio greater than one in all fishing scenarios (Fig. 6, small red circles). This effect was less dramatic with the slot limit, as the slot only protected the female population; i.e., males did not grow beyond the upper size limit (Fig. 1b). For this reason, the shape parameter *p*, representing the degree to which the largest males contributed disproportionately to total care capacity, did not affect the results in Fig. 6.

**Figure 6.**
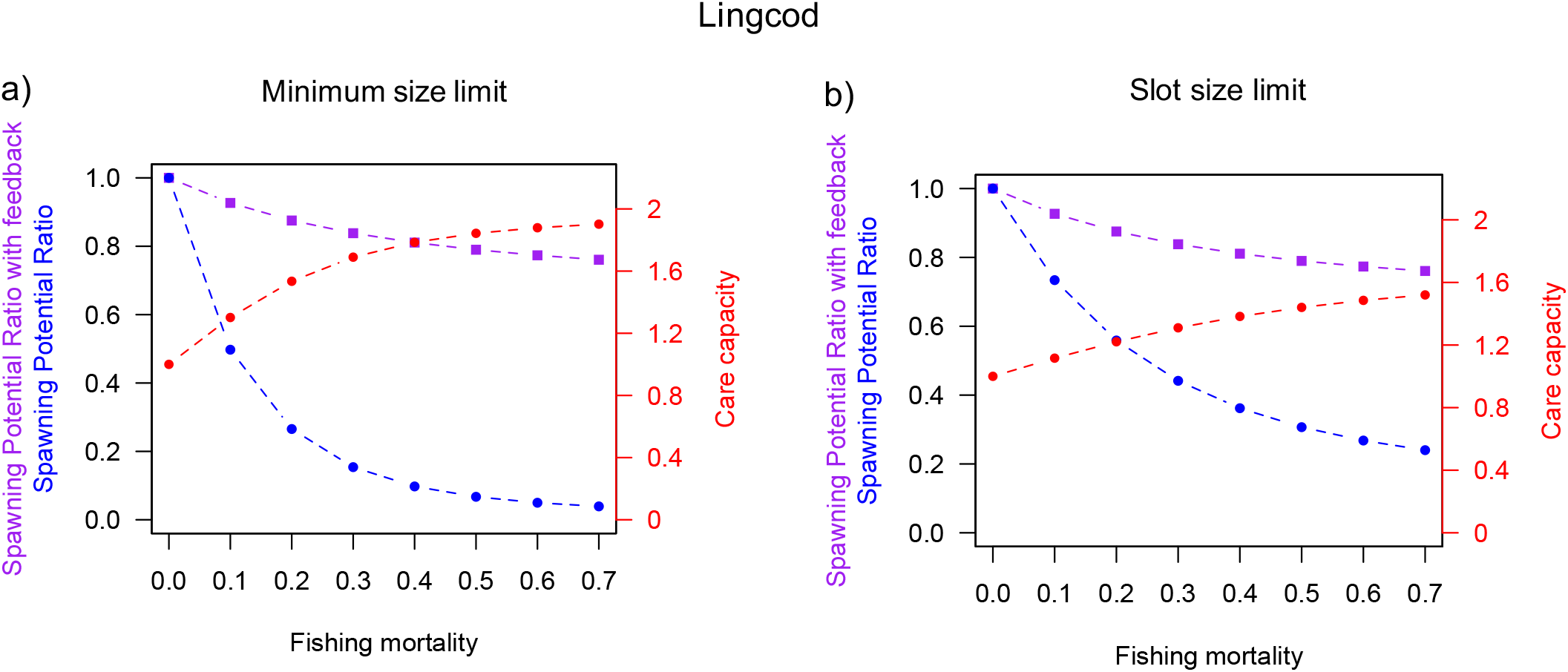
Spawning potential and care capacity for each of the factorial combinations of lingcod simulations. We modeled spawning potential of fisheries with a minimum size limit (left panel) or a slot limit (panel a), each with (squares b) and without (blue-dashed circles) a feedback between care and larval production, across a range of fishing mortalities. Red-dashed circles represent the Care Ratio, R. For lingcod, R values were greater than 1, so care ratio was discounted as it increased (note the right y-axis scale differs from Fig. 4).

## Discussion

Our study is part of a growing body of literature evaluating the potential for size-selective management strategies to balance yield, spawning potential, and long-term fishery sustainability (Gwinn et al., 2015; Kindsvater et al., 2017; Stubberud et al., 2019; Ahrens et al., 2020). The influence of male size structure on population productivity in fished species with obligate male care has been recognized as an important aspect of effective management, but left un-explored (Halvorsen et al., 2016). Here, we modelled fisheries for two such species to investigate the effects of different size-selective management strategies on yield, spawning potential, and care. By using a simulation approach, we were able to investigate the possible importance of an indirect effect (feedback) of fishing on per-egg care, which depends on both male and female size structure. We found that for both species, without any assumptions about how male care influences larval production, the slot limit was most effective at protecting larval production when fishing mortality was low. Under high fishing mortality, in the steady state, very few individuals survived beyond the maximum size limit. This result might change if fishing decreases natural mortality, which is possible if mechanisms of adult mortality are density dependent. This is a plausible mechanism for compensatory productivity, especially for species with high natural mortality (Andersen, Jacobsen, Jansen, & Beyer, 2017; Rose, Cowan, Winemiller, Myers, & Hilborn, 2001). However, our model indicates that slot limits alone could be insufficient to prevent overexploitation if fishing effort overall is not controlled.

In both species, implementing a slot limit led to a difference in yield of 5% or less once fishing mortality (*F*) exceeded 0.4 (Fig. 3b, Fig. 5b). For fishing mortality of this intensity (*F* > 0.4), spawning potential was reduced by at least 60% when parental care affected egg survival in the wrasse. As a whole, our results suggest the growth patterns and mating systems of both species may increase their vulnerability to overfishing, for different reasons, and despite management measures.

### Corkwing wrasse

The benefit of the slot limit to larval production – relative to populations managed with a minimum size limit – could only be detected in the corkwing wrasse simulations when care was reduced by fishing in a way that reduced the proportion of surviving larvae by the fraction *R*. This is because, under fishing, no females grew large enough to benefit from the maximum size limit, so the slot did not affect egg production. A narrower slot that would also provide some protection for large females under low to moderate fishing mortality would likely positively affect both egg supply and care compared to current suggestion of a 120-170 mm slot size limit. The details regarding how care affects larval survival are important. First, we assumed per-capita egg survival decreased linearly with nest availability, i.e. there were no non-linear effects (such as increased survival at low density). This assumption may not hold for all species with obligate male care, but in *S. ocellatus*, a congeneric wrasse species, male care is largely sharable (i.e., we don’t expect a decrease in egg survival at high densities) and at low egg densities, males are more likely to abandon nests (Alonzo 2004). Second, the more nonlinear the relationship between the Male Size Index and care capacity, the more effective the slot limit was at increasing larval production. We do not know how much more effective larger males are in providing care, but in the corkwing wrasse, large males have larger nests and start nesting earlier, which could allow them to care for more eggs and complete more nesting cycles in a spawning season, suggesting that they do provide an outsized contribution to care (Ingebrigt, Uglem & Rosenqvist, 2002).

Large body size of nesting males is likely also favored by sexual selection arising from competition among males and female choice. In our model, size-selective fishing of corkwing wrasse has two impacts: disproportionate removal of nesting males, and decreasing the amount of care available for eggs. We did not model the effect of male removal on fertilization success, because this species also has sneaker males capable of fertilizing eggs, so when some nesting males are present, sperm limitation is unlikely. However, in the long term, any increase in the mating success of these smaller males may be offset by changes in the inter- and intra-sexual dynamics of males and females, as the fitness of sneaker males also depends on the frequency of nesting males and the choosiness of females. Further, if fishing reduces nesting male densities and hence competition over nest territories, it is possible that males can respond by building nests and reproducing at a smaller size (Halvorsen et al., 2016). This behavioral response could compensate for the loss of large nesting males under high fishing intensities, but the smaller body size of nesting males would also imply reduced care. Nevertheless, we chose to not incorporate such a relationship in our model due to the limited empirical knowledge on how population density affects male life history in this and related wrasse species.

### Lingcod

In contrast to the wrasse example, for lingcod, the slot limit was effective in increasing larval production (relative to that expected under a minimum size limit), even without a feedback between nest availability and care capacity (compare Figs. 6a and 6b). However, the fishery selectively removed a greater proportion of the female spawning population than the male population. As a result, this species remained vulnerable to recruitment overfishing, even with a slot limit in place, if total fishing mortality was not controlled. This result is likely to be the case for any species where females are much larger than males and fishing is strongly size-selective (regardless of whether they have male care). While sexual selection on males can favor larger males, the relative strength of fecundity selection on females likely determines whether or not females are larger than males (as in lingcod). Our assumptions about the potential indirect effects of fishing on male care were less important than the direct effect of fishing on egg production. This means that while fishing out larger males could also decrease care-per-egg, for the model scenarios we considered, the effect of fishing on female size structure was more important. For lingcod, even with a slot limit, spawning potential was reduced to 30% or less of unfished levels once fishing mortality exceeded 0.4, unless we assumed there was a “rescue” effect due to the increase in availability of per-egg care (Fig. 6). Currently we do not know whether increased care availability has any such compensatory effect, or if it does, whether the magnitude we considered is realistic. However, our result suggests the use of bag limits, in addition to slot limits, have been important in allowing lingcod stocks to rebuild.

### Broader implications

In our model, recruitment depended on a deterministic Beverton-Holt relationship between larval production and survival. Therefore, our model results regarding yield were not sensitive to our assumptions about feedbacks from male care (Fig. 3; Fig. 5), despite differences in larval production (SPR) arising under different management and harvest scenarios (Fig 4, Fig. 6). In reality, we expect that the relationship between larval supply and yield will fluctuate according to environmental conditions, though detecting this relationship is notoriously tricky (Munch et al., 2018). By using the Beverton-Holt relationship, we assumed there was no overcompensatory density-dependence (as there would be if we used a Ricker function). We chose the Beverton-Holt because it is a more conservative model of recruitment dynamics (i.e., fishing does not increase population productivity as it would under a Ricker model). We also have no evidence for cannibalism at high population densities, or for interference in egg survival at high adult population densities, which have been suggested to drive Ricker-like dynamics in some species. If we had used a Ricker function, the relative increase in larval production expected after implementing a slot size limit would be offset by the decline in recruitment success at high larval densities.

When individuals of different sizes compete for mating success, as in species with alternative reproductive types that differ in size, it is unclear how adaptation to size-selective fishing will affect the size distribution and population productivity (DeFilippo et al., 2019; Kendall et al., 2014; Kendall & Quinn, 2013). Some theory predicts that sexual selection will reinforce adaptation to fishing when males are under directional sexual selection, but fishing increases variance in male reproductive success (Hutchings & Rowe, 2008). There is some evidence from sockeye salmon, however, that the removal of large males by fishing elevates the relative fitness of secondary males (in salmon, known as “jacks”), and increases the frequencies of smaller males in subsequent generations (DeFilippo et al., 2019). In other species with multiple mating tactics, this may not be the case. For example, in a wrasse with paternal care, high sneaking rates have been found to decrease the willingness of nesting males and females to mate (Alonzo & Heckman, 2010; Alonzo & Warner, 1999). In this case, it is possible that sexual selection will actually decrease the success of the remaining (unfished) nesting males in the short term, because the density of small sneaker males at remaining nests will increase. The frequency-dependent selection that is maintaining the stable polymorphism in male mating tactic may be disrupted, and the potential interactions with fishing-induced selection on maturation and growth is unknown. One possibility is that by increasing sneaker density at nests, fishing could indirectly hamper population productivity by decreasing the number of eggs that are spawned, even if females are not fished directly – an undesirable outcome of size-selective management.

In species that do not have secondary males, such as lingcod, it is possible that fertilization rates (sperm limitation) could be a limiting factor in fisheries that target large males (Alonzo and Mangel 2004). However, we found that egg limitation was much more of a concern in our model. Similarly, in the wrasse, the secondary males are assumed to be the outcome of frequency-dependent selection on male mating tactics; if nesting males become rare enough that sperm limitation is an issue. However, over generations, the frequency of each mating type will presumably respond so that more individuals mature as nesting males. For these reasons, we chose not to focus on the effects of sperm limitation in our analyses.

Several studies support the assumption of a positive correlation between male body size and reproductive success in fishes with male care (Bose et al., 2018; Cargnelli & Neff, 2006; Wiegmann & Baylis, 1995). Among fisheries targeting fishes with male care, some are managed by harvest slots, reflecting that the importance of large males (and females) is acknowledged (Table 1). Several of these species are also protogynous hermaphrodites. For these species, management that protects the largest individuals may be necessary to prevent depletion of terminal males to avoid sperm limitation, and can also help to buffer fisheries-induced reduction in size- at sex-change (Alonzo *et al*., 2008; Easter & White, 2016; Easter *et al*., 2020; Kindsvater, Reynolds, Sadovy de Mitcheson, & Mangel, 2017; Sato et al., 2018).

Although harvest slots may seem to be an intuitive measure to balance fishing mortality between sexes, we show that management protecting large individuals of any sex is likely to have small effects if natural mortality is relatively high. That does not mean that large males should be ignored, but rather that management strategies are carefully evaluated and effects monitored. Our results are consistent with three rules of thumb to promote the sustainability of fisheries for species with paternal care: (i) control fishing mortality (e.g. implement quotas or bag limits); (ii) allow both males and females to spawn at least once before fished (e.g. enforce minimum size limits or gear modifications) and (iii) reduce or restrict fishing during the nesting period which can directly affect productivity if guarding males are fished (Froese, 2004; Overzee & Rijinsdorp, 2015; Suski 2003). A spawning season closure has been implemented in the Norwegian wrasse fishery in recent years, but wrasse can be fished during the spawning season in Sweden (Faust, Halvorsen, Andersen, Knutsen, & André, 2018). Lastly, in order to better predict the consequences of harvesting fish with male care, dedicated field studies are needed under realistic conditions. For example, how does disproportional removal of large males affect competition and social hierarchies among remaining males, and to what extent does reduced availability of nests affect female mating decisions and offspring survival?

In summary, our results highlight the need to exert caution when managing fishes with obligate male care, and that slot size limits are no silver bullet for ensuring long term viability if overall fishing mortality is high and not controlled. Furthermore, the evolutionary outcomes of size-selective fishing for the relative fitness of the alternative reproductive tactics deserves further scientific investigation. Despite these caveats, our results suggest that the current management strategies used for both corkwing wrasse in Norway and lingcod in western North America, have likely benefited the sustainability and rebuilding rates of the respective fisheries.

## Acknowledgements

We thank JW White and an anonymous reviewer for comments on a prior version of this manuscript. This work was supported by NSF DEB-1556779 to HKK, funding from Virginia Tech to HKK, NSF IOS-1655297 to SHA, funding from the University of California Santa Cruz to SHA and funding from the Institute of Marine Research (Project 15638-01) to KTH and TKS.

## Data Availability Statement

All data and code analysed in this manuscript are in an archived open-access repository (Kindsvater 2020)

